# Dynamics of a Hes1-Dll1 regulatory network in coupled muscle stem cells: stability, bifurcations, and coexistence of oscillatory states

**DOI:** 10.64898/2026.07.29.741250

**Authors:** Zsófia Bujtár, Björn Goldenbogen, Jana Wolf

## Abstract

Muscle regeneration relies on the coordinated activation of muscle stem cells, whose fate decisions are regulated by intracellular gene expression dynamics and intercellular coupling via the Notch-Dll1 signaling pathway. Central components of this regulatory network include the transcriptional repressor Hes1, its target gene Dll1, and the myogenic regulator MyoD. Experimental and theoretical studies have shown that proliferating muscle stem cells exhibit oscillatory dynamics of these molecules, whereas sustained expression is associated with differentiation. Here, we investigate the dynamics of a previously established delay differential equation model of two coupled muscle stem cells. Using linear stability analysis, we systematically characterize how model parameters affect the transition between stable and unstable steady states. In addition, numerical bifurcation analysis is employed to study the influence of intercellular coupling strength and delay on the system dynamics. Our analysis shows that continuous variation of the coupling delay induces repetitive changes in the stability of the steady state. However, this sensitivity towards the coupling delay is confined to a narrow region of parameter space and therefore requires a fine tuning of all other parameters. Beside the identification of parameter sets for in-phase and out-of-phase oscillations, we demonstrate the possibility of coexisting stable in-phase and out-of-phase oscillations, a dynamical feature that has not been reported previously. While oscillation periods are largely determined by intracellular regulatory mechanisms, oscillation amplitudes can be strongly modulated by intercellular coupling. These results provide new insight into how the intracellular network and intercellular communication interact to generate different collective dynamics.

## 1 Introduction

Mammalian skeletal muscles retain the ability for growth and regeneration throughout life. Muscle regeneration is essential for repair following injury, and impairments in this process occur in genetic diseases, e.g. Duchenne muscular dystrophy, (Guiraud et al. 2019; Ganassi et al. 2022; Deprez et al. 2023) and aging (Blau et al. 2015). Muscle regeneration requires muscle stem cells, which reside as so-called satellite cells between the basal lamina and sarcolemma of myofibers (Musarò 2014). In response to injury, these quiescent cells become activated and proliferate as myoblasts until they differentiate into myocytes to finally fuse and form myofibers (Musarò 2014).

This complex process is tightly regulated by a network of molecular signaling pathways (Musarò 2014) that is also involved in cell fate decisions in other tissues, such as neural stem cells or presomitic mesoderm (Kageyama et al. 2026). Importantly, these regulatory networks often operate through changes in the temporal dynamics of key signaling molecules (Lahmann et al. 2021; Kageyama et al. 2023; Kageyama et al. 2026). Among the signaling pathways involved, the Notch signaling pathway facilitates direct cell-to-cell communication through the coupling of neighboring cells. Notch signaling is activated when the Delta like 1 (Dll1) ligand presented by a signal-sending cell binds to a Notch receptor on an adjacent signal-receiving cell, thereby inducing the expression of transcription factors of the Hes family (Artavanis-Tsakonas et al. 1999; Mumm et al. 2000; Iso et al. 2003; Bray 2006).

Members of the Hes protein family have been found to be key regulators in embryogenesis (Kageyama et al. 2007), e.g. in neurogenesis (Hu et al. 2022) or in hematopoietic stem cell development (Guiu et al. 2013). Common to most of the Hes proteins, such as Hes1, Hes5 and Hes7, is the repressing function on their target genes including their own genes (Kobayashi et al. 2014). The autoinhibition creates a delayed negative feedback which can give rise to oscillatory expression of Hes proteins (Kobayashi et al. 2014). In mice, the expression of Hes proteins oscillates with a period of approximately 2-3 hours, as observed for Hes1 in cultured cells (Hirata et al. 2002) and in muscle stem cells (Lahmann et al. 2019), for Hes5 in neural progenitors (Manning et al. 2019) and for Hes7 in presomitic mesoderm (Bessho et al. 2001).

Hes1 also directly inhibits the transcription of Dll1 (Kobayashi et al. 2009), resulting in an intercellular feedback loop between Hes1 molecules in adjacent cells, as Dll1 activates Notch signaling in these cells. This intercellular feedback via Dll1 can lead to a phase locking between the Hes oscillations of neighboring cells. Hes1-controlled Dll1 protein expression has been observed to oscillate out-of-phase in coupled muscle stem cells (Zhang et al. 2021) and neural progenitors (Shimojo et al. 2016). In contrast, in the presomitic mesoderm, Dll1 and Hes7 expression oscillates in-phase in neighboring cells (Bessho et al. 2001; Shimojo et al. 2016).

In muscle stem cells, Hes1 protein also represses the transcription of *MyoD*, which encodes a skeletal muscle-specific transcription factor that is essential for the differentiation of these cells (Lahmann et al. 2019). It has been shown that an oscillatory dynamics of MyoD is associated with an undifferentiated, proliferating state, required for the maintenance of muscle stem cells, whereas sustained MyoD protein expression is associated with differentiating cells (Lahmann et al. 2019; Lahmann et al. 2021).

Since oscillatory gene expression emerges from regulatory interactions and is modulated by cell-cell coupling, mathematical models have become an important tool for investigating the mechanisms governing Hes dynamics in different cell types. A first ordinary differential equation (ODE) model captured the delayed negative feedback regulation of Hes1 and demonstrated its ability to generate sustained oscillations (Hirata et al. 2002). Other studies used delay differential equations (DDEs) to describe the characteristic transcriptional and translational delays underlying the Hes1 protein oscillation (Jensen et al. 2003; Monk 2003). Models including detailed consideration of the Hes1 regulatory network were used to study the regulation of oscillation amplitudes (Momiji et al. 2008). For muscle stem cells, a model representing the core negative feedback of Hes1 (Hirata et al. 2002) was extended by the myogenic regulator MyoD (Lahmann et al. 2019) and Dll1 (Zhang et al. 2021). To investigate the intercellular communication via the Dll1-Notch signaling pathway and its impact on Hes dynamics, models of two and more cells have been formulated (Lewis 2003; Momiji et al. 2009; Shimojo et al. 2016; Yoshioka-Kobayashi et al. 2020; Zhang et al. 2021; Giri et al. 2021; Biga et al. 2021; Hawley et al. 2022; Stevens et al. 2026). The Notch signaling pathway includes several processes such as receptor-ligand interactions, cleavage of the Notch intracellular domain and its passage to the nucleus, which define the strength and the delay of the intercellular coupling. Variation of the coupling delay can change the phase relation of oscillations and can induce oscillation death (Momiji et al. 2009; Shimojo et al. 2016; Yoshioka-Kobayashi et al. 2020; Zhang et al. 2021; Kageyama et al. 2023; Kageyama et al. 2026). However, for a given coupling delay, either in-phase or out-of-phase oscillations were found (Shimojo et al. 2016; Yoshioka-Kobayashi et al. 2020; Zhang et al. 2021; Kageyama et al. 2023).

Here, we revisit the two muscle stem cell model proposed by Zhang et al. (Zhang et al. 2021) to systematically investigate the impact of all model processes on the system dynamics. In a stability analysis, we characterized the influence of all model parameters on the transition between stable and unstable steady states.

We further employ bifurcation analysis to investigate the impact of the intercellular coupling on the dynamics. By varying coupling strength and delay, we could identify, in contrast to previous studies, parameter regimes in which in-phase and out-of-phase periodic orbits coexist. Furthermore, we show that while the oscillation period is largely determined by intracellular regulatory processes, the amplitudes of the oscillations can be modulated by the characteristics of intercellular coupling.

### 1.1 The two-coupled-cell model

This study focuses on a published model for the dynamics of the Hes1-Dll1 gene regulatory network in two mouse muscle stem cells coupled via Dll1-Notch signaling (Zhang et al. 2021). The qualitative model, schematically shown in Figure 1a, represents the simplified signaling network. It describes the dynamics of Hes1 and Dll1 protein concentrations in both cells, denoted by *H*_1_ and *H*_2_ for Hes1 and *D*_1_ and *D*_2_ for Dll1, respectively (Eqs. 1-4). The model includes the autoinhibition of Hes1 protein (Hes1p), the intracellular inhibition of Dll1 protein (Dll1p) expression by Hes1p, as well as the intercellular activation of Hes1p expression by Dll1p. Through the negative regulation of Dll1p expression by Hes1p and the positive regulation of Dll1p on Hes1p in the neighboring cell, a double negative regulation loop, i.e. positive feedback loop, is formed. In this model, processes including multiple subsequent events such as transcription, translation and transportation are lumped into combined reactions with increased time requirement incorporated by time delays (*τ*). Here, *τ*_1_ and *τ*_21_ represent the time required for Hes1p to affect the concentrations of Hes1p and Dll1p in the same cell, respectively, while *τ*_22_ represents the time required for Dll1p to affect the Hes1p concentration in the neighboring cell. In particular, *τ*_21_ captures the transcriptional delay of Dll1p expression, and *τ*_1_ and *τ*_22_ include the transcriptional delay of Hes1p expression. Since the sum of *τ*_21_ and *τ*_22_ represents the overall delay between Hes1p or Dll1p in the two coupled cells, we will refer to it as ‘total coupling delay’ in the following.

**Fig. 1.**
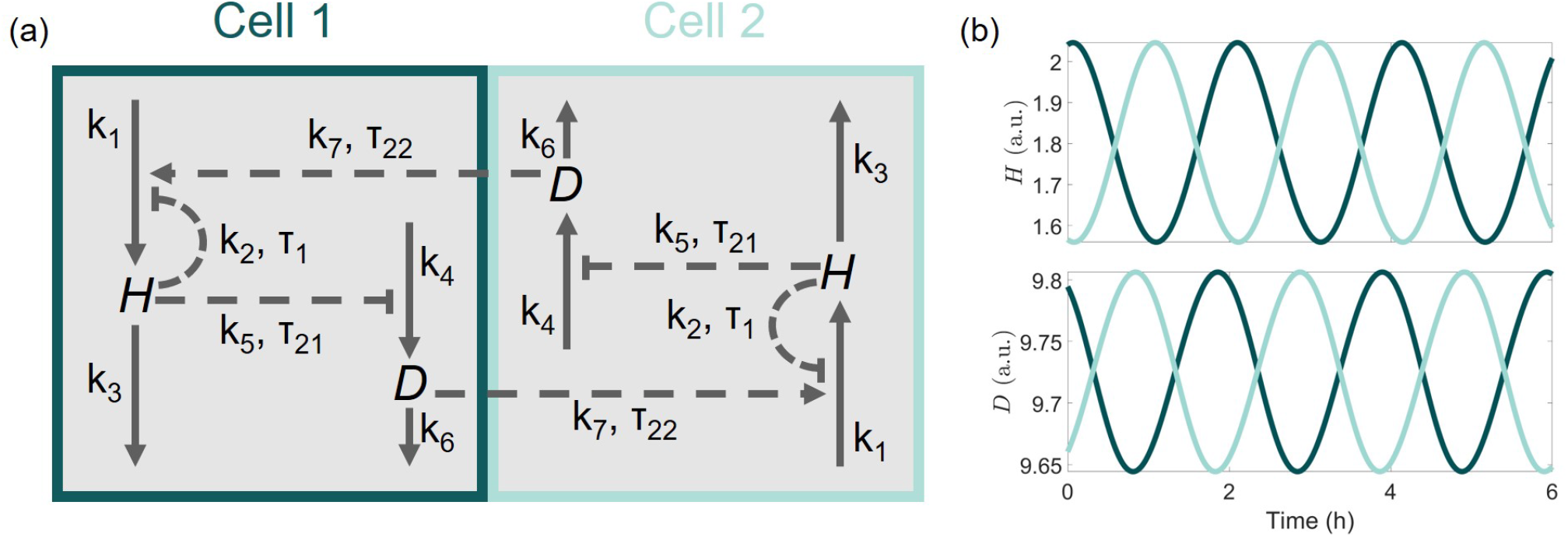
Description of the two-coupled-cell model by Zhang et al. (Zhang et al. 2021). (a) Model scheme showing the regulation of protein concentrations of Hes1 (*H*) and Dll1 (*D*) in the two cells (cell 1 shown in dark green, cell 2 in light green). Production of both proteins is inhibited by Hes1p within the cell. Dll1p activates production of Hes1p in the neighboring cell. The inhibitory regulations are characterized by the delays *τ*_1_ and *τ*_21_, the activation by delay *τ*_22_. Processes are shown including the relevant rate constants (*k*_1_, …, *k*_7_) and delays. (b) Out-of-phase oscillations at reference parameter set, parameter values are given in Supplementary Table 1.

The degradation of both Hes1p and Dll1p is described by first-order kinetics with rate coefficients *k*_3_ and *k*_6_, respectively, whose values are based on experimental data (Hirata et al. 2002; Shimojo et al. 2016). The production of Hes1p is induced by the Dll1p concentration of the neighboring cell delayed by *τ*_22_ with the scaling rate coefficient 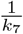 and the rate constant *k*_1_. The production is inhibited by Hes1p in the same cell delayed by *τ*_1_. The synthesis of Dll1p occurs constitutively with rate constant *k*_4_ and is inhibited by Hes1p with the delay *τ*_21_. The strength of the inhibition by Hes1p on itself and Dll1p is modulated by the parameters *k*_2_ and *k*_5_, respectively, which refer to the delayed concentrations of Hes1p, for which the inhibition effect is half maximal. In this form, two, independent Hes1 DNA-bindings are considered at the two regulatory mechanisms of Hes1p.

In the following we consider the model of two identical cells, i.e. both cells are described by the same model structure and the same parameter set.

An exemplary model simulation with reference parameter values (Supplementary Table 1) is shown in Figure 1b. Under this condition, *H* and *D* in both cells exhibit an oscillatory dynamics with a period of around 2 hours and an out-of-phase relation between the two coupled cells (see details of simulations in Section 4.1). It reproduces the experimental observation for activated, proliferative wild-type muscle stem cells very well (Zhang et al. 2021).

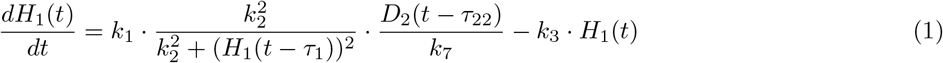

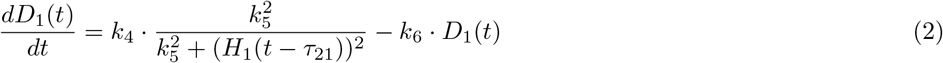

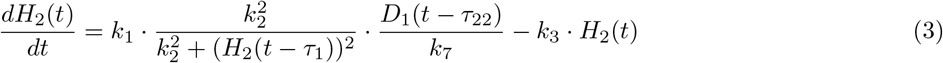

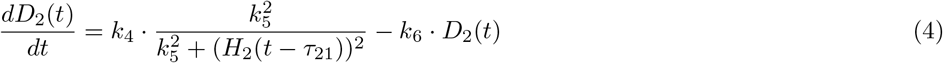

## 2 Results

### 2.1 Changing the delay parameters *τ*_21_ and *τ*_22_ leads to repetitive stability changes

As oscillatory and sustained expression of the key transcription factor Hes1 is associated with proliferation and differentiation of muscle stem cells, respectively, we wanted to identify conditions of oscillatory and sustained dynamics based on our mathematical representation of the system. Therefore, we calculated the steady states with respect to the parameter set and determined their local stability based on the eigenvalues of the characteristic equation of the linearized system at those steady states, using toolbox DDE-BIFTOOL (Engelborghs et al. 2001; Sieber et al. 2014) (see details in Section 4.2). The parameters *k*_1_, …, *k*_7_ were varied between fold change (FC) of 0.1 and 4.0 w.r.t. the reference parameter set, while the delays *τ*_1_, *τ*_21_, *τ*_22_ were varied between 0 h and 20 h.

In the DDE model, the steady states are, by definition, independent of the value of any time delay parameter, so their values can only depend on the parameters *k*_1_, …, *k*_7_. The steady states were numerically estimated for the reference parameter set and individual variations of the parameters *k*_1_, …, *k*_7_. For the reference parameter set, a steady state with real values was identified, where both cells have the same equilibrium and by varying any parameter *k*_1_, …, *k*_7_ individually, the steady state concentrations gradually change. The steady state values of *H* and *D* for the different parameter sets are shown in Supplementary Figure 1. Here, alterations of the two basal production rate constants *k*_1_ and *k*_4_, have a similar effect on the steady state concentration of *H*. Also, alterations of the two degradation rate coefficients *k*_3_, *k*_6_ as well as coupling-related parameter *k*_7_, impact the steady state concentration of *H* equally. Comparing the impact of the parameters on the steady state of *H* and *D*, those parameters that describe the Hes1-related mechanisms (*k*_1_, *k*_2_, *k*_3_, *k*_7_) have different effects from those involved in the Dll1-related mechanisms (*k*_4_, *k*_5_, *k*_6_). Changing any of the Dll1-related parameters, the steady state of the two protein concentrations changes in a similar direction. By contrast, the Hes1-related parameters have the opposite effect on the steady state of *D* than on the steady state of *H*: at increased Hes1 parameter, the steady state of either *D* or *H* is increased while the other is decreased.

For the identified steady states, their stability with respect to the parameter set has been assessed. In contrast to the steady state itself, its stability is not independent of the time delays, therefore also the impact of the delay parameters on the stability had to be considered. The stability of a steady state was determined based on its eigenvalues of the linearized system at this steady state. If there are no eigenvalues with positive real part, the steady state is stable, if there exists at least one, the steady state is unstable. Since for DDEs the number of eigenvalues is infinite, only the eigenvalues with the highest real part are shown for all individual parameter modifications in Figure 2. The delays are shown in absolute values, while changes in all other parameters are represented as fold changes w.r.t. to the reference set (Supplementary Table 1). The parameters can be divided into four groups (a)-(d) based on the pattern of their impact on the eigenvalues in the studied parameter range (1^*st*^ column, Figure 2). For groups (a) and (b), a stable and an unstable steady-state region were identified. The first group (a) comprises the parameters *k*_1_, *k*_4_ and *τ*_1_, for which the stable steady state changes to the unstable steady state when the parameter is increased. The direction of the change is reversed for the second group (b), comprising parameters *k*_2_, *k*_6_ and *k*_7_. For parameters *k*_3_ and *k*_5_, which form the third group (c), there are stable steady-state regions for high and low parameter values and an unstable steady-state region in between. In the fourth group (d), encompassing parameters *τ*_21_ and *τ*_22_, the real parts of eigenvalues successively change their sign for increasing parameters, leading to repetitive changes between stable and unstable steady states. Comparing the effect of the different parameters on the steady state stability, we conclude that the parameters *k*_1_ and *k*_7_ have inverse impacts, as their product is invariant. The two basal production rate coefficients *k*_1_ and *k*_4_ have similar impacts, i.e. leading to stable steady state at lower coefficients and unstable steady state at higher ones. Comparing the two degradation rate coefficients *k*_3_ and *k*_6_ with each other or the delay *τ*_1_ with *τ*_21_ or *τ*_22_, their impacts on the stability are qualitatively different.

**Fig. 2.**
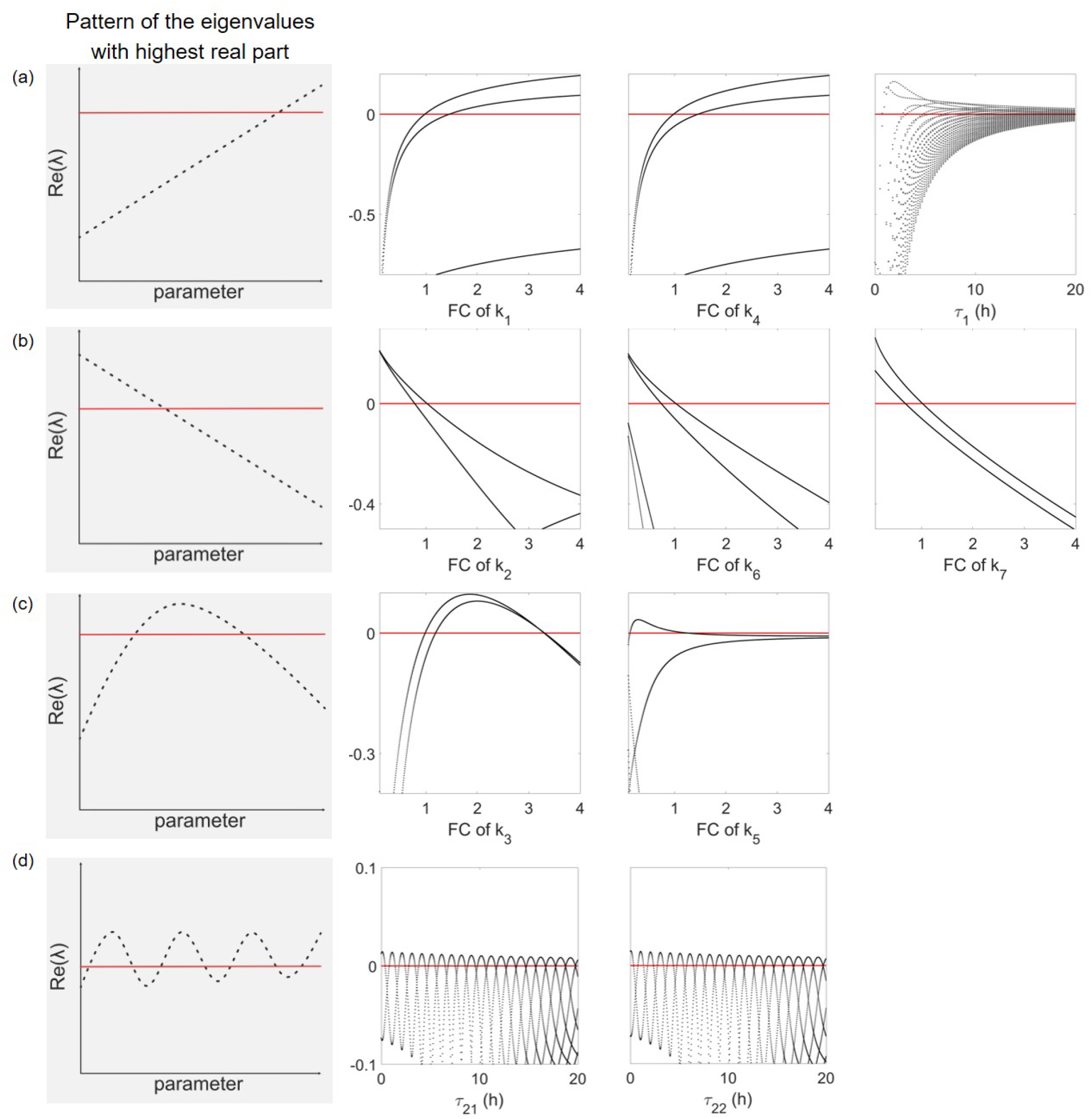
Parameter variation causes distinct patterns of eigenvalues with highest real. Increase in parameter values lead to different pattern of the real parts of the eigenvalues, which can cross the zero line (shown in red), and therefore stability changes (a) from stable to unstable, (b) from unstable to stable, (c) from stable to unstable and then back to stable, or (d) repetitively between unstable and stable. Common trends of the eigenvalues with highest real part for each group of parameters is visualized in first column. *k*_1_, …, *k*_7_ were varied from 0.1 FC to 4.0 FC w.r.t the corresponding reference values, while *τ*_1_, *τ*_21_, *τ*_22_ were varied between 0 and 20 h. Only eigenvalues with the highest real parts are shown.

Based on the observation that individual changes in *τ*_21_ and *τ*_22_ in the reference parameter set lead to characteristic repetitive stability changes, we explored the questions how general this observation is and how other parameters impact this stability relation.

### 2.2 Impact of the total coupling delay on the steady state stability

To compare the effect of parameters *τ*_21_ and *τ*_22_ on steady state stability, both parameters were varied independently in the range from 0 to 6 h (Figure 3). The analysis shows that the steady state stability depends on the value of both parameters equally, as indicated by a diagonal symmetry in the *τ*_21_ - *τ*_22_ parameter space, i.e. changes of *τ*_22_ can be compensated by changes of *τ*_21_ of the same magnitude. While this observation is based on the numerical estimation of the eigenvalues of the characteristic equation of the linearized system at the found steady state and fixed parameters *k*_1_, …, *k*_7_, *τ*_1_, it can be generalized by an analytical calculation of the characteristic equation of the linearized system (see details in Section 4.3). The two delay parameters *τ*_21_ and *τ*_22_ can be replaced by their sum, *τ*_21_ + *τ*_22_, so that the eigenvalues of the characteristic equation only depend on the sum of the two delays as a single parameter. This sum or total coupling delay corresponds to the time required for Hes1p or Dll1p in one cell to react on the change of Hes1p or Dll1p in the neighboring cell.

**Fig. 3.**
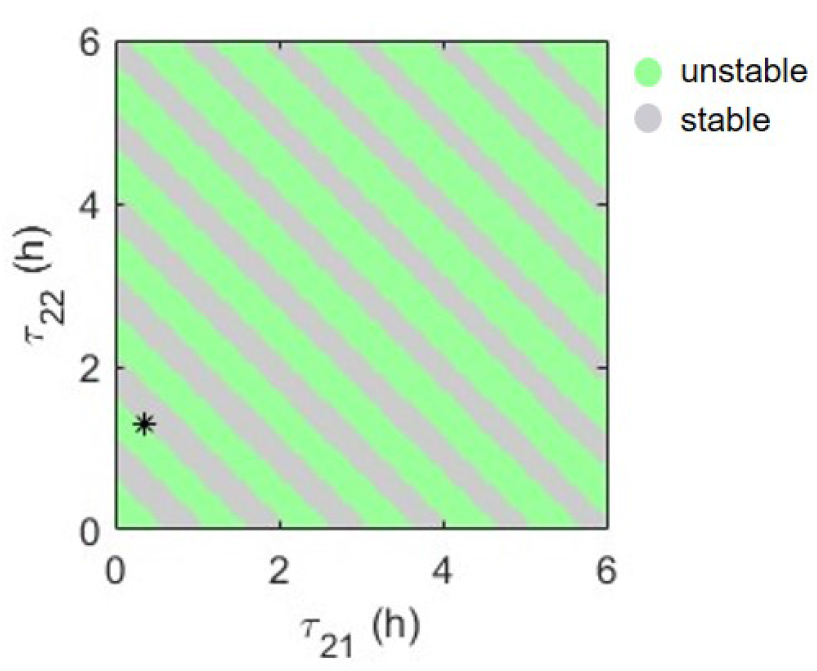
Stability of the steady state is invariant to changes in delays *τ*_21_ and *τ*_22_ that preserve their sum. Shown are stable and unstable steady state regions for varying delays *τ*_21_ and *τ*_22_ in range from 0 to 6 h. Point of reference parameter values (*τ*_21_ = 0.35 h, *τ*_22_ = 1.3 h) is marked by a star.

Having identified a repetitive change in the steady state stability for a constantly increasing or decreasing total coupling delay, we investigated the contribution of the other kinetic parameters on this stability pattern. To this end, we investigated combinations of the total coupling delay and one of the other parameters *k*_1_, …, *k*_7_ or *τ*_1_. For each parameter combination, the stability of the steady state was determined and plotted in stability maps (Figure 4).

**Fig. 4.**
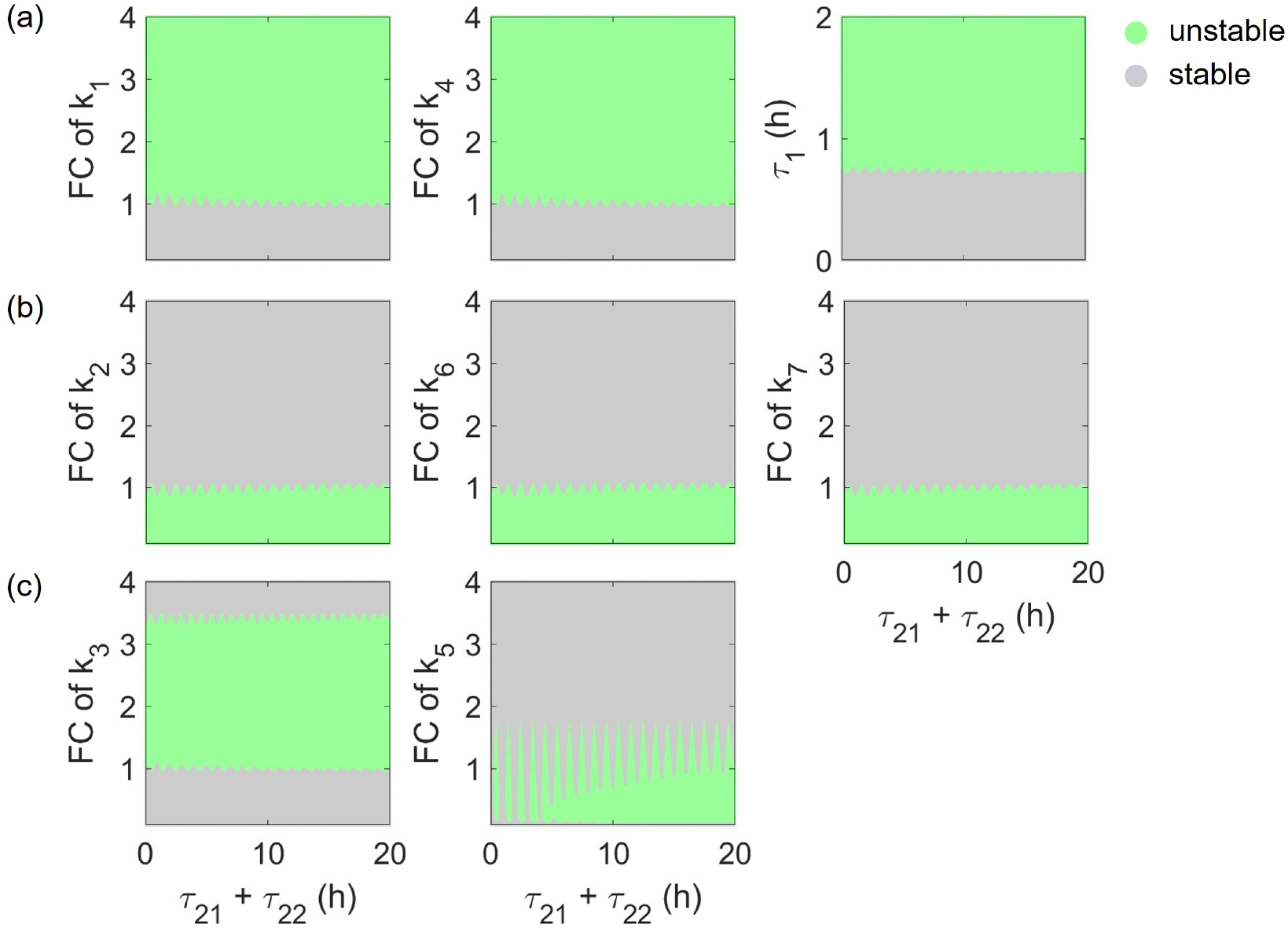
Combined impact of the total coupling delay *τ*_21_ + *τ*_22_ and the parameter k_1_, …, k_7_, *τ*_1_ on the steady state stability. Shown are stability maps as a function of parameters *k*_1_, …, *k*_7_ or *τ*_1_ and total coupling delay *τ*_21_ + *τ*_22_. (a) An increase in *k*_1_, *k*_4_ or *τ*_1_ leads to unstable steady states, (b) while an increase in *k*_2_, *k*_6_ or *k*_7_ leads to stable steady states. The boundary between the stable and unstable steady-state regions shows a wave-like pattern along the axis of the total coupling delay. (c) For *k*_3_, an increase leads first to unstable, then to stable steady state. A complex stability pattern is observed for the combination of *k*_5_ and *τ*_21_ + *τ*_22_ with islands of unstable steady states for lower values of *τ*_21_ + *τ*_22_. For higher values of *τ*_21_ + *τ*_22_, an increase in parameter *k*_5_ can lead first to unstable, then back to stable steady state, similarly to *k*_3_. The kinetic parameters *k*_1_, …, *k*_7_ were varied individually from 0.1 FC to 4 FC, and *τ*_1_ was varied between 0 and 2 h. The total coupling delay (*τ*_21_ + *τ*_22_) was varied in the range from 0 to 20 hours including the reference value 1.65 h.

The stability pattern along the horizontal lines at FC = 1 for parameters *k*_1_, …, *k*_7_ and at *τ*_1_ = 0.725 h corresponds to the effect of varying the total coupling delay (*τ*_21_ + *τ*_22_) while keeping all other parameters at their reference values. Therefore, it shows the repetitive change in the stability described above. Vertical changes at the reference value of the total coupling delay, *τ*_21_ + *τ*_22_ = 1.65 h, correspond to parameter changes in Figure 2a-c. Increasing the parameters *k*_1_, *k*_4_, or *τ*_1_ lead to an unstable steady state (Figure 4a), whilst increasing the parameters *k*_2_, *k*_6_, or *k*_7_ lead to a stable steady state (Figure 4b). Subfigures 4a and 4b show that these trends remain consistent over a broad range of the total coupling delay. In addition, these subfigures reveal that over a wide parameter range of *k*_1_, *k*_4_, *τ*_1_, *k*_2_, *k*_6_ and *k*_7_, the stability of the steady state is independent of the total coupling delay and there are only small regions in which the total coupling delay impacts the stability of the steady state. In contrast to the afore mentioned parameters, *k*_3_, the degradation constant of Hes1p, shows three distinct stability regions, in the investigated parameter range. Increasing *k*_3_ from 0 first leads to a stability transition at around 1 FC from stable to unstable. However, when increasing *k*_3_ even further to values *>* 3.5 FC, the stability of the steady state switches back to stable. Note that, in the vicinity of both transition-points of *k*_3_, changes in the total coupling delay will lead to the described repetitive switching from stable to unstable to stable (Figure 4c left).

The stability pattern shows the most differences for the parameter *k*_5_. Here, a repetitive change in stability for *τ*_21_ + *τ*_22_ variation occurs in a wider range of *k*_5_. Noteworthy, for low values of the total coupling delay, distinctive islands with unstable steady states emerge in the *k*_5_ –(*tau*_21_ + *tau*_22_) - plane. While at the islands of the unstable steady states, oscillatory dynamics can occur (Supplementary Figure 3), left and right column), stable steady-state regions are characterized by sustained dynamics independent of the *k*_5_ in FC (see Supplementary Figure 3, middle column). See also Supplementary Figure 2 for the real parts of the eigenvalues for *k*_5_-FC-variation. At higher FC values of *k*_5_, only stable steady states occur (exemplary simulation shown in Supplementary Figure 3 (upper line)), whereas combinations of low FC values of *k*_5_ and higher total coupling delay, the islands vanishes and a connected region with unstable steady state emerges.

These analyses demonstrate that the total coupling delay can alter the stability of the steady state only within certain narrow ranges of all other kinetic parameters. A quantification of these individual ranges for all parameters at the reference parameter set is given at Supplementary Table 2, again demonstrating that this total coupling delay sensitive range is bigger for *k*_5_ than for *k*_1_, *k*_2_, *k*_3_, *k*_4_, *k*_6_, *k*_7_ w.r.t. to their fold changes. This narrow parameter range that defines the total coupling delay sensitive region, implies that a fine tuning of the kinetic parameters is required to allow for an impact of the coupling delay on the qualitative system dynamics. The reference parameter set is positioned in the region, where changes in delay parameters *τ*_21_ and *τ*_22_ can change the dynamics, which is highly relevant for the prediction of mutant behavior.

### 2.3 The effect of a Dll1 transcriptional delay on the dynamics depends on the actual setting of the system

The network dynamics in cells with mutations resulting in prolonged Dll1 transcriptional delay has been investigated (Shimojo et al. 2016; Yoshioka-Kobayashi et al. 2020; Zhang et al. 2021). Experiments and simulations for muscle stem cells showed that a 0.1-hour prolongation of the transcriptional delay of Dll1 (*τ*_21_) shifts the dynamics from oscillations to sustained expression (Zhang et al. 2021), see Figure 5, lower left and right. In another mutant scenario, the effect of the prolongation of the Hes1 transcriptional delay (impacting *τ*_1_ and *τ*_22_ here) by around 0.1 h was studied (Ochi et al. 2020) and they experimentally and computationally observed a slightly prolonged oscillations period of Hes1p. We here use the two-coupled-cell model to predict the effect of a combined mutation, i. e. prolonged *τ*_1_, *τ*_22_ and *τ*_21_. The time required for Hes1p expression is included in the delay of Hes1 autoinhibition described by parameter *τ*_1_ as well as in the delay of Hes1 activation by Dll1p described by parameter *τ*_22_, therefore both of these parameters were increased simultaneously to simulate mutant cells with prolonged Hes1 transcriptional delay (see details of simulations in Section 4.1). The simulations show that a simultaneous increase in *τ*_1_ and *τ*_22_ by 0.1 h leads to oscillations here with higher amplitude of *H* and slightly increased period (Figure 5, lower left and upper left). A combination of this mutational effect with an alteration increasing the Dll1 transcriptional delay (*τ*_21_) by 0.1 h predicts that in this combined mutational case the cells do not go to stable steady state but keep oscillating (Figure 5, upper right). However, this prediction of a conditions specific effect of a *τ*_21_-increase remains to be experimentally tested.

**Fig. 5.**
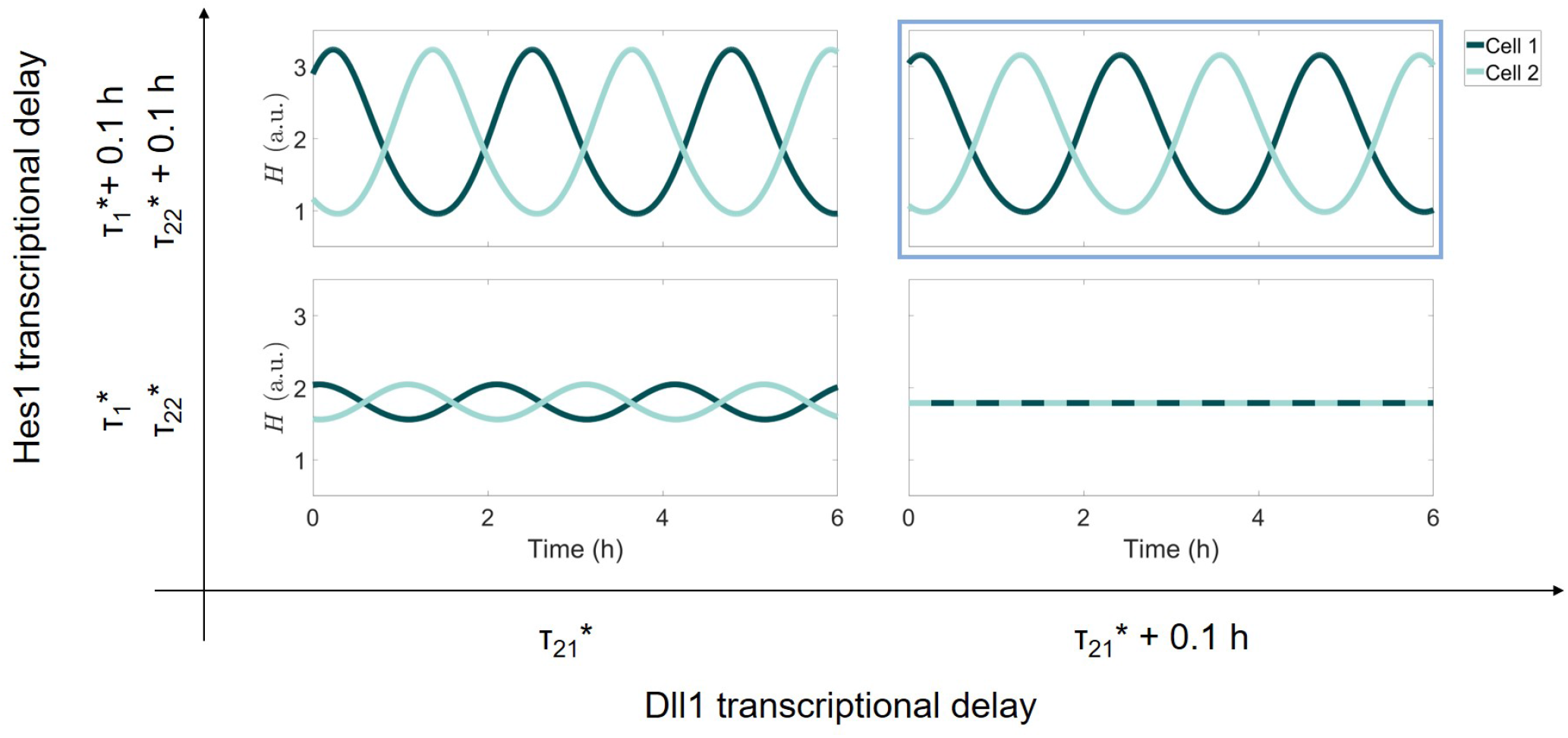
Dependence of oscillatory dynamics on the Dll1 transcriptional delay at reference versus increased Hes1 transcriptional delay. It was predicted that the oscillatory dynamics of wild-type conditions (reference parameter set) vanishes if the Dll1 transcriptional delay (*τ*_21_) is prolonged by 0.1 h (Zhang et al. 2021) (lower left and right panels). For an increase in the Hes1 transcriptional delay (increasing *τ*_1_ and *τ*_22_ by 0.1 h), oscillations can be observed (upper left panel). We here predict that an additional increase of the Dll1 transcriptional delay (*τ*_21_) by 0.1 h in this case does not abolish oscillations (upper right panel outlined by a blue box). Reference parameter values are shown by *\**.

### 2.4 Intercellular coupling strength and delay strongly impact the phase difference and Hes1 amplitude of periodic orbits

Our detailed analysis allows to investigate the impact of the intercellular coupling on the cellular Hes1 dynamics. Specifically, we focus on the effect of changes in intercellular coupling parameters: the delay *τ*_22_ and the parameter *k*_7_ that is inversely proportional to the coupling strength.

Linear stability analysis provides only a local information on the system dynamics in the vicinity of the steady state. Therefore, it remained unanswered what dynamics would be observed in the unstable steady-state region and under what coupling conditions periodic solutions could arise. In particular, we were interested in the region around the reference parameter set, as an oscillatory dynamics for *H* and *D* has been already reported for this parameter set. We systematically analyzed the system dynamic in the interval between 1.05 h and 2.05 h for *τ*_22_ and between 0.6 FC and 1.04 FC for *k*_7_, thereby including the reference parameter values (Figure 6, Supplementary Figure 4). Note that the interval for the total coupling delay was varied between 1.4 h and 2.4 h, since *τ*_21_ = 0.35 h in the reference set. Please refer to Figure 4 for stability analysis with respect to the total coupling delay and intercellular coupling strength.

**Fig. 6.**
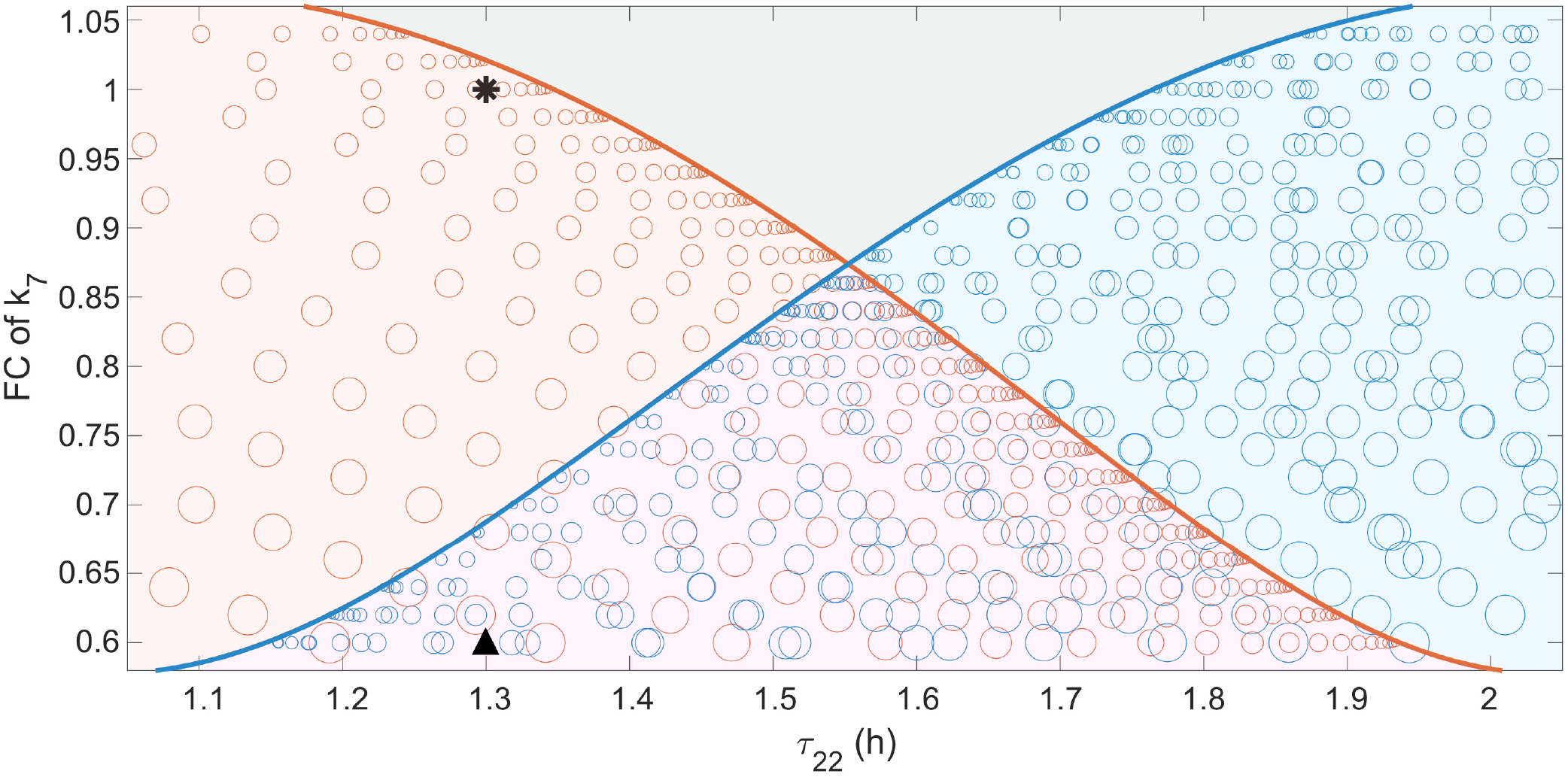
Identification of subregions with oscillatory and non-oscillatory dynamics for different values of intercellular coupling delay and strength. Subregions are identified based on the number of eigenvalues with positive real parts: zero (gray), two (red and blue) or four (purple). Two bifurcation curves (shown in red and blue) separate the subregions and intersect at the codimension-2 double Hopf bifurcation point. The gray subregion refers to a stable steady state, while the other three subregions refer to out-of-phase (red), in-phase (blue) and coexisting in-phase and out-of-phase periodic orbits (purple) around an unstable steady state. The area of the circles is proportional to the amplitude of *H* at the periodic orbits. Point of reference (*τ*_22_ = 1.3 h, FC of *k*_7_ = 1) is marked by a star. The triangle marks the parameter pair (*k*_7_, *τ*_22_), for which two simulations leading to in-phase and out-of-phase oscillations are shown in Figure 7.

In the analyzed two-dimensional parameter space of *τ*_22_ and *k*_7_, the unstable steady-state region is separated from the stable steady-state region by bifurcation lines: as the imaginary part of the eigenvalues with highest real part are non-zero, these are Hopf bifurcations. The unstable steady-state region itself is divided in three subregions, colored in red, blue and purple in Figure 6. In the red and blue regions, there are two eigenvalues with positive real part, while in the purple region, there are four of those eigenvalues. In the stable steady-state region, colored in gray in Figure 6, there is no eigenvalue with positive real part. The regions of stable and unstable steady states are separated by two bifurcation lines intersecting at the codimension-2 Hopf-Hopf (or double Hopf) bifurcation point (Kuznetsov 2004).

Within the three unstable steady-state subregions, periodic orbits were identified, using the toolbox DDE-BIFTOOL (Engelborghs et al. 2001; Sieber et al. 2014) (see details in Section 4.2). We characterized these periodic orbits with respect to phase difference between the coupled cells and and with respect to the amplitude and period of *H*. We found in-phase and out-of-phase periodic orbits, with phase differences of 0 and half a period between *H* in the two cells, respectively. On the left side of the red curve, we observed out-of-phase periodic orbits, while on the right side of the blue curve, we observed in-phase periodic orbits. This implies that for a specific FC value of *k*_7_ out-of-phase and in-phase oscillations can occur for different *τ*_22_ values. For example, for *k*_7_ at 0.6 FC, low *τ*_22_ values lead to out-of-phase periodic orbits and high *τ*_22_ values lead to in-phase periodic orbits. Interestingly, at the middle region of *τ*_22_, marked in purple in Figure 6, we found coexisting in-phase and out-of-phase periodic orbits. Exemplary simulations starting from different initial conditions are shown at Figure 7, where *τ*_22_ kept its reference value (details in Section 4.1). We also quantified the amplitude and the period of the found periodic orbits for the studied two-coupled-cell model (Figure 6 and Supplementary Figure 4). The amplitude of *H* periodic orbits (shown by the area of the circles at Figure 6) is very small close to Hopf bifurcation points, but increases with increasing distance from the bifurcation line. This opens the possibility that coexisting in-phase and out-out-phase periodic orbits have similar or different amplitudes, depending on the parameter set and their distances from the bifurcation lines. The oscillation period is almost constant at 2 h in the analyzed region, for both, the in-phase and out-of-phase periodic orbits (Supplementary Figure 4).

**Fig. 7.**
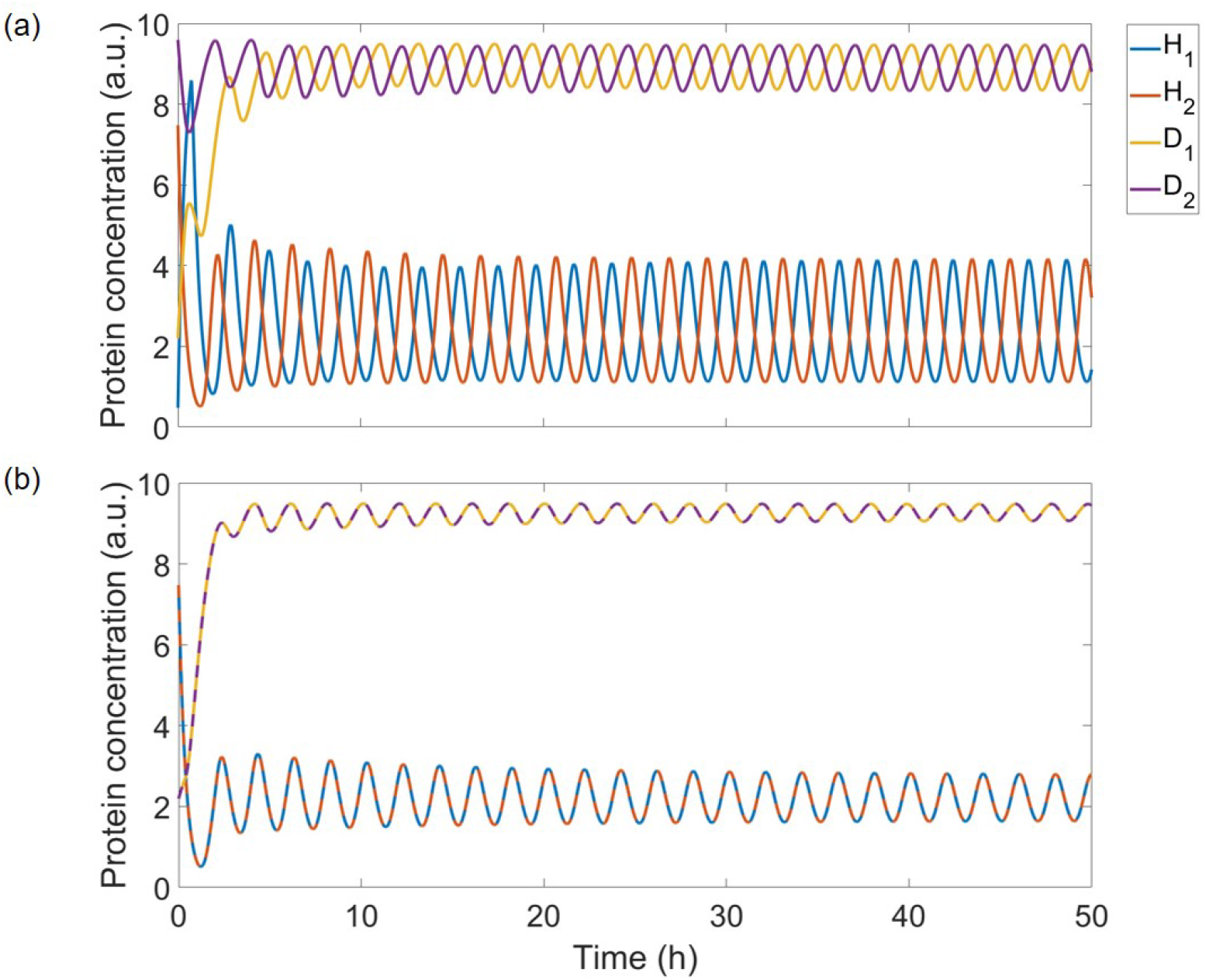
Coexisting (a) out-of-phase and (b) in-phase oscillations. The simulations show different trajectories for different initial conditions but same parameterization: *k*_7_ = 0.6 FC and *τ*_22_ = 1.3 h. Remaining parameters were set to the reference values.

## 3 Discussion

Following a muscle injury, proliferation and differentiation of muscle stem cells must be balanced to ensure optimal tissue regeneration. The corresponding cell fate decisions depend on the dynamics of MyoD, which in turn is controlled by a Hes1 regulatory network including intercellular coupling processes. Using a model of two coupled cells (Zhang et al. 2021), we investigated the influence of the parameters of Hes1-related processes and intercellular coupling on signal dynamics.

Stability analysis of the model showed that all kinetic parameters, including those involved in syntheses, degradations and intracellular regulations, impact the dynamical behavior of the coupled-cell model. For a specific parameter range around the reference parameter set, we could confirm the described effect that variations of the coupling delay can shift the system dynamics from oscillatory to sustained and vice versa (Momiji et al. 2009; Shimojo et al. 2016; Yoshioka-Kobayashi et al. 2020; Zhang et al. 2021). However, outside of this parameter range, variations of the coupling delay are insufficient to change the stability of the steady state. Of note, even relatively small and biologically plausible changes in parameters can drive the system out of the delay-sensitive region, e.g. by a fold change of 1.2 in the synthesis rates *k*_1_ or *k*_4_, or the degradation rate *k*_6_. However, since the effect of the coupling delay on the system dynamics has been observed experimentally, our finding implies a fine-tuned interplay of cellular and intercellular processes in wild type cells that allow for correct systems dynamics and resulting cell-fate decisions. In our study, the parameter *k*_5_, characterizing the inhibitory regulation of Hes1 on Dll1 within a cell, displayed the broadest range of the parameter fold change in which the stability of the steady state was sensitive for the total coupling delay. The relevance of the sensitivity of the system dynamics towards the coupling delay is demonstrated by investigating a double mutant effect. Model simulations predict that the experimentally and computationally well-characterized amplitude death observed for a prolonged Dll1 transcriptional delay (*τ*_21_ + 0.1 h) in muscle stem cells (Zhang et al. 2021) can be recovered by a prolongation of the Hes1 transcriptional delay (*τ*_1_ + 0.1 h and *τ*_22_ + 0.1 h). Our results show that the reported sensitivity of cell fate decisions towards delay in transcription of Dll1 and the total coupling delay could be restricted to a rather small range of the overall parameter space. The demonstrated impact of multiple cellular processes on the dynamics shows the sensitivity of the system towards all rates of the network and might be explored in the future to see how alterations of other processes, e.g. by pharmacological interventions, might perturb the dynamics involved in muscle stem cell function and muscle regeneration (Deprez et al. 2023).

The stability analysis showed that the delays *τ*_21_ (Dll1 transcriptional delay) and *τ*_22_ (intercellular coupling delay) had a similar effect on the steady state stability and can be lumped to a total coupling delay (*τ*_21_ + *τ*_22_). Continuously increasing or decreasing this coupling delay leads to repetitive switching between stable and unstable steady states, a phenomenon that has also been observed in a related two-cell model (Momiji et al. 2009) and other systems, e.g. a damped oscillator with delayed friction or delayed restoring force (Cooke et al. 1982) or two simple oscillators coupled with delay (Ramana Reddy et al. 1998).

To further investigate the impact of the intercellular coupling strength (1*/k*_7_) and delay (*τ*_22_), we performed bifurcation analyses of the system around the reference parameter set, thereby identifying a double Hopf bifurcation point at the intersections of two bifurcation curves. Such double Hopf bifurcation point has been already reported for other systems (Li et al. 2012; Song et al. 2022).

Each of the two bifurcation curves is associated with the amplitude death of one of the stable periodic orbits for Hes1 in the coupled cells, although with different phase shifts. Hence, the two bifurcation curves separate the parameter space into four subregions near the double Hopf bifurcation point, in which the system dynamics differs significantly. Only in one subregion the steady state is stable. In two of the subregions, the Hes1 concentration of the coupled cells oscillates either exclusively in-phase or exclusively out-of-phase, similar to reports from other studies (Shimojo et al. 2016; Yoshioka-Kobayashi et al. 2020; Zhang et al. 2021; Kageyama et al. 2023). However, in one subregion we observed the coexistence between stable in-phase and out-of-phase oscillations. In that subregion, the system can reach one of the coexisting periodic orbits depending on the initial condition. The amplitudes of coexisting in-phase and out-of-phase periodic orbits are either very similar or different, depending on the distance to the corresponding bifurcation curve, and therefore on the specific values of the intercellular coupling and strength. However, the oscillatory periods were found to be very similar and rather independent of the intercellular coupling strength and delay, so they are mainly determined by intracellular regulatory mechanisms. The possible coexistence of in-phase and out-of-phase oscillations indicates, that coupled Hes oscillators do not always have to follow a strict phase-relation. Such multi-rhythmicity has been observed for other biological oscillators, as reviewed in (Goldbeter et al. 2022; Tyson et al. 2022). It remains to be explored in how far this occurrence of bistability, described before in a more general form (Reddy et al. 2000; Pfeuty 2022), can contribute to the observed microclusters of coupled cells that are characterized by synchrony in otherwise out-of-phase oscillating cell populations (Biga et al. 2021; Hawley et al. 2022).

Our focus on the model of two interacting and identical cells allowed the systematic analyses of all parameter effects on the dynamics of the Hes1. Extending this investigation to models of larger and more heterogeneous cell populations for muscle stem cells remains an open task. However, earlier studies in other cell lines demonstrated, that results established in 2-coupled-cell models are predictive also for larger cell populations (Shimojo et al. 2016; Yoshioka-Kobayashi et al. 2020; Kageyama et al. 2023), and additional modes of dynamical behavior can arise from inclusion of heterogeneity of cells or their coupling mechanisms (Dey et al. 2021; Hawley et al. 2022; Ho et al. 2024).

## 4 Methods

### 4.1 Model simulations

A built-in MATLAB function, dde23, was used to integrate the delay differential equation system that approximates the solution numerically. As delay differential equations require the history of the solution until the highest time delay in the past as initial condition, the history of the solution was set to constant, more specifically to non-negative and real values, as *H* and *D* refer to protein concentrations.

### 4.2 Local stability analysis and periodic orbit identification

For both, assessing the local stability for the steady state and identification of periodic orbits around an unstable steady state, the MATLAB toolbox DDE-BIFTOOL (Engelborghs et al. 2001; Sieber et al. 2014) was used.

The steady state was numerically estimated for a defined parameter set and its stability was determined based on the eigenvalues of the characteristic equation. The eigenvalues are approximated using a numerical, linear multistep method. After calculating the initial steady state and its corresponding eigenvalues for a defined parameter set, a branch of steady states was calculated by varying a parameter of interest in a defined interval. Along these parameter sets, branches of respective eigenvalues were computed. Although for DDEs, the set Λ = {*λ* : *λ* ∈ ℂ}, the number of eigenvalues corresponding to a steady state at a defined parameter set, is infinite, the set *S* = {Re(*λ*) : Re(*λ*) *>* 0, *λ ∈* Λ}, the number of eigenvalues in any right half plane, is not, therefore, the stability is always determined by a finite number of eigenvalues (Sieber et al. 2014). Knowing the eigenvalues with the highest real part for a given steady state at a defined parameter set, let us determine its stability. If *S* = *∅*, i.e. there exists no eigenvalue with positive real part, the steady state is stable and vice versa if *S /*= *∅*, i.e. there exists minimum one eigenvalue with positive real part, the steady state is unstable. Note that if an eigenvalue crosses the imaginary axes, it refers to a bifurcation point. If the max({Re(*λ*) : *λ* ∈ Λ}) = 0 and the corresponding imaginary part of this eigenvalue is non-zero, Im(*λ*) ≠ 0, it is called Hopf bifurcation point.

From a bifurcation point, where an eigenvalue crosses the imaginary axes at a non-zero imaginary part, a branch of periodic solutions can emanate. After finding the bifurcation point at a given parameter set, along an interval of a chosen parameter, a branch of periodic orbits was identified. For the found periodic orbits, the phase difference between variables, the amplitude and the period were quantified. The phase difference between the coupled cells and the amplitude with respect to the intercellular coupling are shown in Figure 6, where the MATLAB built-in scatter function was used to visualize the periodic orbit with circle: its color refers to in-phase or out-of-phase periodic orbit and its area is proportional to the oscillatory amplitude of *H*. The period with respect to the intercellular coupling is presented at Supplementary Figure 4.

### 4.3 Characteristic equation of the linearized DDE model

Numerical calculation showed that changes in the sum of the delays *τ*_21_ and *τ*_22_ can change the stability of the steady state. However, concurrent changes of *τ*_21_ and *τ*_22_ that do not change the sum left the stability of the steady state unchanged. This dependency of the stability of a steady state on kinetic parameters can be shown using the characteristic equation of the linearized system.

The characteristic equation of a general linearized system of DDEs

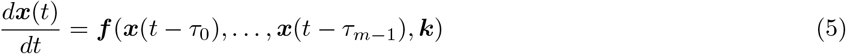

with *m* constant delays and *τ*_0_ = 0 is given by

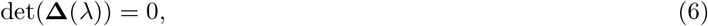

in which the matrix **Δ** *∈* ℝ^*m×m*^ with respect to the eigenvalue *λ* is defined as

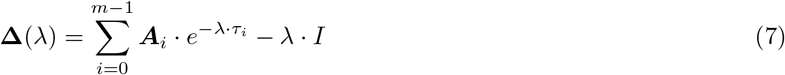

with *I* ∈ ℝ^*m×m*^ being the identity matrix and ***A***_*i*_ ∈ ℝ^*m×m*^ for *i* ∈ {0, *· · ·, m* − 1} being the delay-depending Jacobian matrix defined as

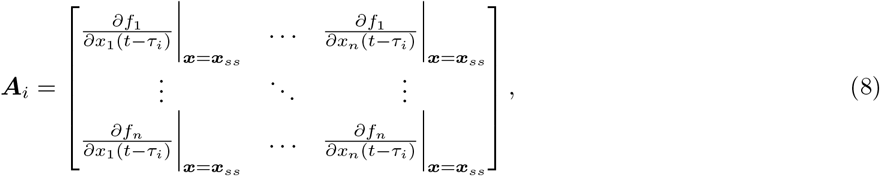

where ***x***_*ss*_ is the steady state, for which ***f*** (***x***_*ss*_) = 0 (Sieber et al. 2014).

For the investigated two-coupled-cell model (Zhang et al. 2021), the Jacobian matrices ***A***_*i*_ *∈* ℝ^4*×*4^ for *i ∈ {*0, 1, 21, 22*}* with the individual delays *τ*_0_, *τ*_1_, *τ*_21_ and *τ*_22_, respectively, are

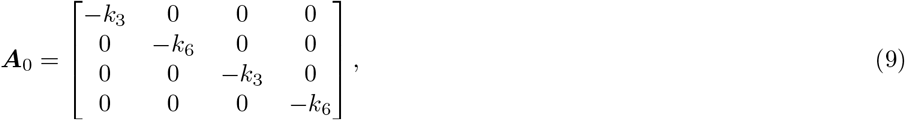

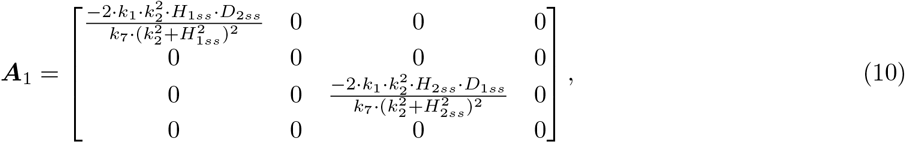

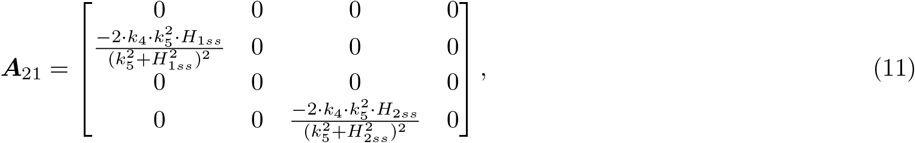

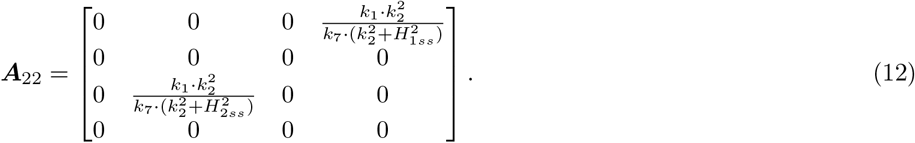

Substituting Eqs. 9-12 into the characteristic equation 6, we obtain the characteristic equation as

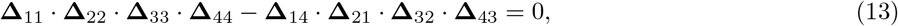

where the matrix elements **Δ**_*ij*_ for *i, j ∈ {*1, *· · ·*, 4*}* are given by

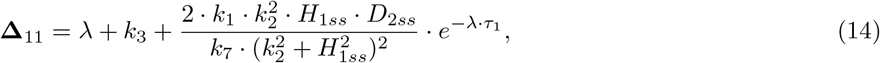

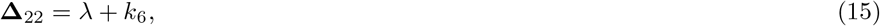

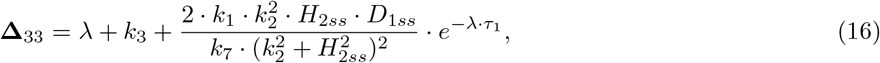

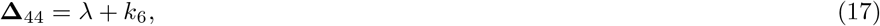

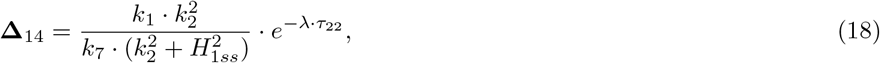

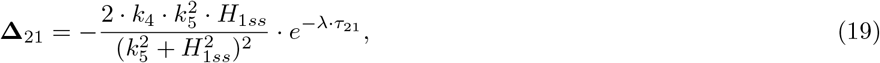

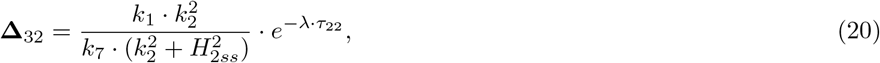

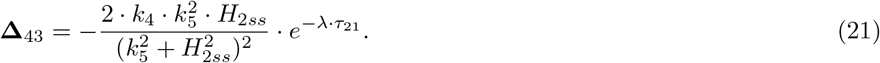

Substituting Eqs. 14-21 into Eq. 13 and rearranging the resulting characteristic equation with respect to terms that are dependent or independent of the delay parameters *τ*_1_, *τ*_21_ and *τ*_22_, we obtain the characteristic equation as

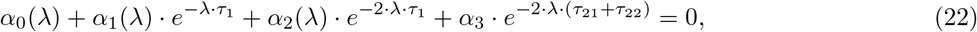

where the left-hand side of the equation is the sum of *τ*-dependent exponential functions with *τ*-independent coefficients *α*_0_, …, *α*_3_. The coefficients *α*_0_, …, *α*_2_ *∈* ℂ might depend additionally to *λ* on the steady state and the parameters. For *α*_3_ *∈* ℝ it holds that *α*_3_ ≤ 0.

From Eq. 22, it can be clearly seen that the delay parameters *τ*_21_ and *τ*_22_ only contribute through their sum to the characteristic equation, as observed at the studied parameter space for the identified steady state through the numerical estimation of eigenvalues.

## Statements and Declarations

The authors declare no competing interests. The authors declare that they have no known competing financial interests or personal relationships that could have appeared to influence the work reported in this paper.

## Acknowledgment

We thank Carmen Birchmeier, Thomas Müller, and Shanshan You for many fruitful discussions and valuable biological insights that helped shape this study. We also thank Sina Glöckner for carefully checking the mathematical derivations, equations, and calculations and Mareike Simon for proofreading.

## Author contributions

**J.W**. and **B.G**. conceived the study. **Zs.B**. performed the mathematical analyses, implemented the computational methods, carried out the simulations and bifurcation analyses, and generated the figures. **J.W**. and **B.G**. supervised the project. All authors contributed to the interpretation of the results, jointly wrote and revised the manuscript, and approved the final version.

## References

Artavanis-Tsakonas, Spyros, Matthew D. Rand, and Robert J. Lake (1999). “Notch signaling: cell fate control and signal integration in development”. In: Science (New York, N.Y.) 284.5415. 10.1126/SCIENCE.284.5415.770.

Bessho, Yasumasa, Ryoichi Sakata, Suguru Komatsu, Kohei Shiota, Shuichi Yamada, and Ryoichiro Kageyama (2001). “Dynamic expression and essential functions of Hes7 in somite segmentation”. In: Genes & development 15.20. 10.1101/GAD.930601.

Biga, Veronica, Joshua Hawley, Ximena Soto, Emma Johns, Daniel Han, Hayley Bennett, Antony D Adamson, Jochen Kursawe, Paul Glendinning, Cerys S. Manning, and Nancy Papalopulu (2021). “A dynamic, spatially periodic, micro-pattern of HES5 underlies neurogenesis in the mouse spinal cord”. In: Molecular Systems Biology. 10.15252/msb.20209902.

Blau, Helen M., Benjamin D. Cosgrove, and Andrew T.V. Ho (2015). “The central role of muscle stem cells in regenerative failure with aging”. In: Nature medicine 21.8. 10.1038/NM.3918.

Bray, Sarah J. (2006). “Notch signalling: a simple pathway becomes complex”. In: Nature Reviews Molecular Cell Biology 7.9. 10.1038/nrm2009.

Cooke, Kenneth L and Zvi Grossman (1982). “Discrete Delay, Distributed Delay and Stability Switches”. In: Journal of Mathematical Analysis and Applications 86.2.

Deprez, Alyson, Zakaria Orfi, Lupann Rieger, and Nicolas Alexandre Dumont (2023). “Impaired muscle stem cell function and abnormal myogenesis in acquired myopathies”. In: Bioscience Reports 43.1. 10.1042/BSR20220284/232343.

Dey, Supravat, Lee Tracey, and Abhyudai Singh (2021). “Role of intercellular coupling and delay on the synchronization of genetic oscillators”. In: Proceedings of the American Control Conference 2021-May. 10.23919/ACC50511.2021.9482935.

Engelborghs, Koen, Tatyana Luzyanina, and Giovanni Samaey (2001). DDE-BIFTOOL v. 2.00: a Matlab package for bifurcation analysis of delay differential equations. Tech. rep. KU Leuven. URL: https://www.cs.kuleuven.be/publicaties/rapporten/tw/TW305.pdf.

Ganassi, Massimo, Francesco Muntoni, and Peter S. Zammit (2022). “Defining and identifying satellite cell-opathies within muscular dystrophies and myopathies”. In: Experimental Cell Research 411.1. 10.1016/J.YEXCR.2021.112906.

Giri, Amitava, Dola Sengupta, and Sandip Kar (2021). “Deciphering the Role of Fluctuation Dependent Intercellular Communication in Neural Stem Cell Development”. In: ACS Chemical Neuroscience 12.13. 10.1021/acschemneuro.1c00116.

Goldbeter, Albert and Jie Yan (2022). “Multi-synchronization and other patterns of multi-rhythmicity in oscillatory biological systems”. In: Interface Focus 12.3. 10.1098/RSFS.2021.0089.

Guiraud, Simon and Kay E. Davies (2019). “Regenerative biomarkers for Duchenne muscular dystrophy”. In: Neural Regeneration Research 14.8. 10.4103/1673-5374.253534.

Guiu, Jordi, Ritsuko Shimizu, Teresa D’Altri, Stuart T. Fraser, Jun Hatakeyama, Emery H. Bresnick, Ryoichiro Kageyama, Elaine Dzierzak, Masayuki Yamamoto, Lluis Espinosa, and Anna Bigas (2013). “Hes repressors are essential regulators of hematopoietic stem cell development downstream of Notch signaling”. In: Journal of Experimental Medicine 210.1. 10.1084/JEM.20120993.

Hawley, Joshua, Cerys Manning, Veronica Biga, Paul Glendinning, and Nancy Papalopulu (2022). “Dynamic switching of lateral inhibition spatial patterns”. In: Journal of the Royal Society Interface 19.193. 10.1098/rsif.2022.0339.

Hirata, Hiromi, Shigeki Yoshiura, Toshiyuki Ohtsuka, Yasumasa Bessho, Takahiro Harada, Kenichi Yoshikawa, and Ryoichiro Kageyama (2002). “Oscillatory expression of the BHLH factor Hes1 regulated by a negative feedback loop”. In: Science 298.5594. 10.1126/science.1074560.

Ho, Christine, Laurent Jutras-Dubé, Michael L Zhao, Gregor ID Mönke István Z Kiss, Paul François, and Alexander Aulehla (2024). “Nonreciprocal synchronization in embryonic oscillator ensembles”. In: Proceedings of the National Academy of Sciences of the United States of America. 10.1073/pnas.2401604121.

Hu, Nan and Linqing Zou (2022). “Multiple functions of Hes genes in the proliferation and differentiation of neural stem cells”. In: Annals of Anatomy 239. 10.1016/j.aanat.2021.151848.

Iso, Tatsuya, Larry Kedes, and Yasuo Hamamori (2003). “HES and HERP families: multiple effectors of the Notch signaling pathway”. In: Journal of cellular physiology 194.3. 10.1002/JCP.10208.

Jensen, Mogens H., K. Sneppen, and G. Tiana (2003). “Sustained oscillations and time delays in gene expression of protein Hes1”. In: FEBS Letters 541. 1–3. 10.1016/S0014-5793(03)00279-5.

Kageyama, Ryoichiro and Akihiro Isomura (2026). “Oscillatory Gene Expression During Cell Differentiation”. In: Annual Review of Cell and Developmental Biology 48.7. 10.1146/annurev-cellbio-111524-093438.

Kageyama, Ryoichiro, Akihiro Isomura, and Hiromi Shimojo (2023). “Biological Significance of the Coupling Delay in Synchronized Oscillations”. In: Physiology 38.2. 10.1152/PHYSIOL.00023.2022.

Kageyama, Ryoichiro, Toshiyuki Ohtsuka, and Taeko Kobayashi (2007). “The Hes gene family: repressors and oscillators that orchestrate embryogenesis”. In: Development (Cambridge, England) 134.7. 10.1242/DEV.000786.

Kobayashi, Taeko and Ryoichiro Kageyama (2014). “Expression dynamics and functions of hes factors in development and diseases”. In: Current Topics in Developmental Biology. Vol. 110. Academic Press Inc. 10.1016/B978-0-12-405943-6.00007-5.

Kobayashi, Taeko, Hiroaki Mizuno, Itaru Imayoshi, Chikara Furusawa, Katsuhiko Shirahige, and Ryoichiro Kageyama (2009). “The cyclic gene Hes1 contributes to diverse differentiation responses of embryonic stem cells”. In: Genes and Development 23.16. 10.1101/gad.1823109.

Kuznetsov, Yuri A. (2004). “Numerical Analysis of Bifurcations”. In: Elements of applied bifurcation theory. Springer. URL: https://link.springer.com/chapter/10.1007/978-1-4757-3978-7_10.

Lahmann, Ines, Dominique Bröhl, Tatiana Zyrianova, Akihiro Isomura, Maciej T. Czajkowski, Varun Kapoor, Joscha Griger, Pierre Louis Ruffault, Despoina Mademtzoglou, Peter S. Zammit, Thomas Wunderlich, Simone Spuler, Ralf Kühn, Stephan Preibisch, Jana Wolf, Ryoichiro Kageyama, and Carmen Birchmeier (2019). “Oscillations of MyoD and Hes1 proteins regulate the maintenance of activated muscle stem cells”. In: Genes and Development 33. 9–10. 10.1101/gad.322818.118.

Lahmann, Ines, Yao Zhang, Katharina Baum, Jana Wolf, and Carmen Birchmeier (2021). “An oscillatory network controlling self-renewal of skeletal muscle stem cells”. In: Experimental Cell Research 409.2. 10.1016/j.yexcr.2021.112933.

Lewis, Julian (2003). “Autoinhibition with Transcriptional Delay: A Simple Mechanism for the Zebrafish Somitogenesis Oscillator”. In: Current Biology 13. 10.1016/s0960-9822(03)00534-7.

Li, Yanqiu, Weihua Jiang, and Hongbin Wang (2012). “Double Hopf bifurcation and quasi-periodic attractors in delay-coupled limit cycle oscillators”. In: Journal of Mathematical Analysis and Applications 387.2. 10.1016/j.jmaa.2011.10.023.

Manning, Cerys S., Veronica Biga, James Boyd, Jochen Kursawe, Bodvar Ymisson, David G. Spiller, Christopher M. Sanderson, Tobias Galla, Magnus Rattray, and Nancy Papalopulu (2019). “Quantitative single-cell live imaging links HES5 dynamics with cell-state and fate in murine neurogenesis”. In: Nature Communications 10.1. 10.1038/s41467-019-10734-8.

Momiji, Hiroshi and Nicholas A.M. Monk (2008). “Dissecting the dynamics of the Hes1 genetic oscillator”. In: Journal of Theoretical Biology 254.4. 10.1016/j.jtbi.2008.07.013.

Momiji, Hiroshi and Nicholas A.M. Monk (2009). “Oscillatory Notch-pathway activity in a delay model of neuronal differentiation”. In: Physical Review E - Statistical, Nonlinear, and Soft Matter Physics 80.2. 10.1103/PhysRevE.80.021930.

Monk, Nicholas A.M. (2003). “Oscillatory Expression of Hes1, p53, and NF-kB Driven by Transcriptional Time Delays”. In: Chemistry & Biology 10. 10.1016/s0960-9822(03)00494-9.

Mumm, Jeffrey S. and Raphael Kopan (2000). “Notch Signaling: From the Outside In”. In: Developmental Biology 228.2. 10.1006/DBIO.2000.9960.

Musarò, Antonio (2014). “The Basis of Muscle Regeneration”. In: Advances in Biology 2014.1. 10.1155/2014/612471.

Ochi, Shohei, Yui Imaizumi, Hiromi Shimojo, Hitoshi Miyachi, and Ryoichiro Kageyama (2020). “Oscillatory expression of Hes1 regulates cell proliferation and neuronal differentiation in the embryonic brain”. In: Development (Cambridge) 147.4. 10.1242/dev.182204.

Pfeuty, Benjamin (2022). “Multistability and transitions between spatiotemporal patterns through versatile Notch-Hes signaling”. In: Journal of Theoretical Biology 539. 10.1016/j.jtbi.2022.111060.

Ramana Reddy, D. V., A. Sen, and G. L. Johnston (1998). “Time delay induced death in coupled limit cycle oscillators”. In: Physical Review Letters 80.23. 10.1103/PhysRevLett.80.5109.

Reddy, D V Ramana, A Sen, and G L Johnston (2000). “Experimental Evidence of Time-Delay-Induced Death in Coupled Limit-Cycle Oscillators”. In: Physical Review Letters. 10.1103/PhysRevLett.85.3381.

Shimojo, Hiromi, Akihiro Isomura, Toshiyuki Ohtsuka, Hiroshi Kori, Hitoshi Miyachi, and Ryoichiro Kageyama (2016). “Oscillatory control of Delta-like1 in cell interactions regulates dynamic gene expression and tissue morphogenesis”. In: Genes and Development 30.1. 10.1101/gad.270785.115.

Sieber, Jan, Koen Engelborghs, Tatyana Luzyanina, Giovanni Samaey, and Dirk Roose (2014). “DDE-BIFTOOL Manual - Bifurcation analysis of delay differential equations”. In: 10.48550/arxiv.1406.7144.

Song, Yongli, Yahong Peng, and Tonghua Zhang (2022). “Double Hopf Bifurcation Analysis in the Memory-based Diffusion System”. In: Journal of Dynamics and Differential Equations. 10.1007/s10884-022-10180-z.

Stevens, Angela and Nicola Vassena (2026). Mathematical modeling and analysis of the Notch-Delta pathway. URL: https://arxiv.org/pdf/2604.05888.

Tyson, John J., Attila Csikasz-Nagy, Didier Gonze, Jae Kyoung Kim, Silvia Santos, and Jana Wolf (2022). “Time-keeping and decision-making in living cells: Part II”. In: Interface Focus 12.4. 10.1098/rsfs.2022.0024.

Yoshioka-Kobayashi, Kumiko, Marina Matsumiya, Yusuke Niino, Akihiro Isomura, Hiroshi Kori, Atsushi Miyawaki, and Ryoichiro Kageyama (2020). “Coupling delay controls synchronized oscillation in the segmentation clock”. In: Nature 580.7801. 10.1038/s41586-019-1882-z.

Zhang, Yao, Ines Lahmann, Katharina Baum, Hiromi Shimojo, Philippos Mourikis, Jana Wolf, Ryoichiro Kageyama, and Carmen Birchmeier (2021). “Oscillations of Delta-like1 regulate the balance between differentiation and maintenance of muscle stem cells”. In: Nature Communications 12.1. 10.1038/s41467-021-21631-4.

