## Supplementary Information for "Dynamics of a Hes1-Dll1 regulatory network in coupled muscle stem cells: stability, bifurcations, and coexistence of oscillatory states"

**Journal of Mathematical Biology**

**Zsófia Bujtár**[a,b]**, Björn Goldenbogen**[a]**, Jana Wolf**[a,b*]

a) Max Delbrück Center for Molecular Medicine in the Helmholtz Association, Mathematical Modelling of Cellular Processes, Berlin, Germany

b) Freie Universität Berlin, Department of Mathematics and Computer Science, Berlin, Germany

| Parameter | Reference value | Unit |
|---|---|---|
| $k_1$ | 100.0 | 1/h |
| $k_2$ | 1.0 | - |
| $k_3$ | 2.6 | 1/h |
| $k_4$ | 11.2 | 1/h |
| $k_5$ | 5.5 | - |
| $k_6$ | 1.04 | 1/h |
| $k_7$ | 50.0 | - |
| $\tau_1$ | 0.725 | h |
| $\tau_{21}$ | 0.35 | h |
| $\tau_{22}$ | 1.3 | h |

**Supplementary Table 1 Reference parameter values of the two-coupled-cell model (Zhang et al. 2021).** They are positive and real numbers without or with unit of time or its reciprocal.

| Parameter | min FC | max FC | max/min |
|---|---|---|---|
| $\tau_1$ | 0.98 | 1.05 | 1.07 |
| $k_3$ | 0.96 | 1.08 | 1.13 |
| $k_7$ | 0.88 | 1.06 | 1.20 |
| $k_2$ | 0.89 | 1.08 | 1.21 |
| $k_1$ | 0.94 | 1.15 | 1.22 |
| $k_4$ | 0.93 | 1.14 | 1.23 |
| $k_6$ | 0.88 | 1.09 | 1.24 |
| $k_5$ | 0.14 | 1.79 | 12.79 |

**Supplementary Table 2 Estimated fold change ranges and ratios for $\mathbf{k_1, ..., k_7}$ and $\tau_1$ of the total coupling delay-dependent region around the reference parameter value.**

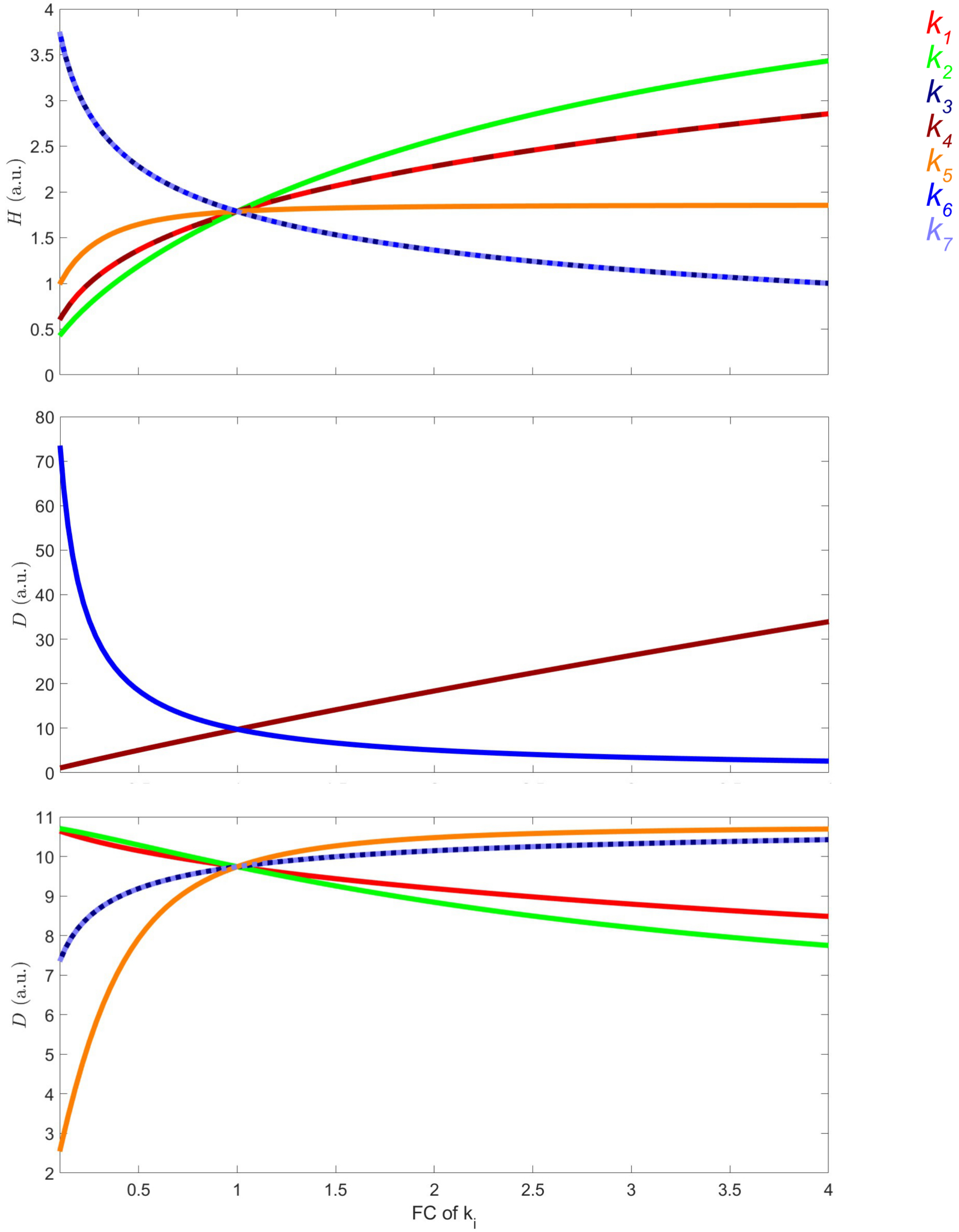


**Supplementary Figure 1 Steady states values of H and D in both cells with respect to parameters $\mathbf{k_1, ..., k_7}$.** Parameters were varied individually between 0.1 FC and 4.0 FC.

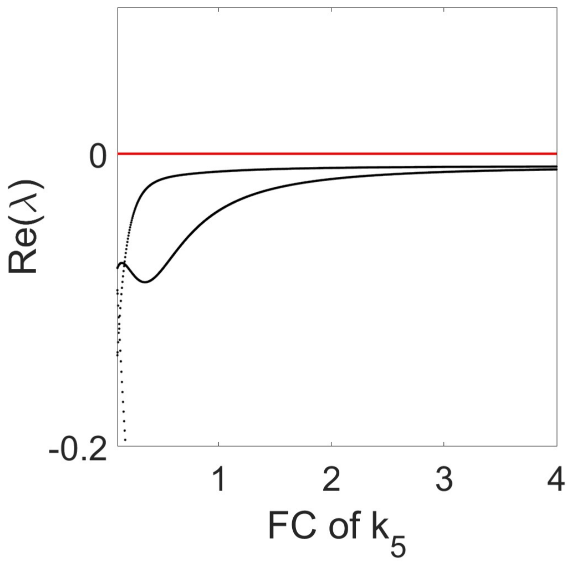


**Supplementary Figure 2 Variation of parameter $\mathbf{k_5}$ in case of a total coupling delay $(\tau_{21} + \tau_{22})$ of 2 h shows only stable steady states.** The eigenvalues with highest real part have only negative real parts. Parameter $k_5$ was varied from 0.1 FC to 4.0 FC.

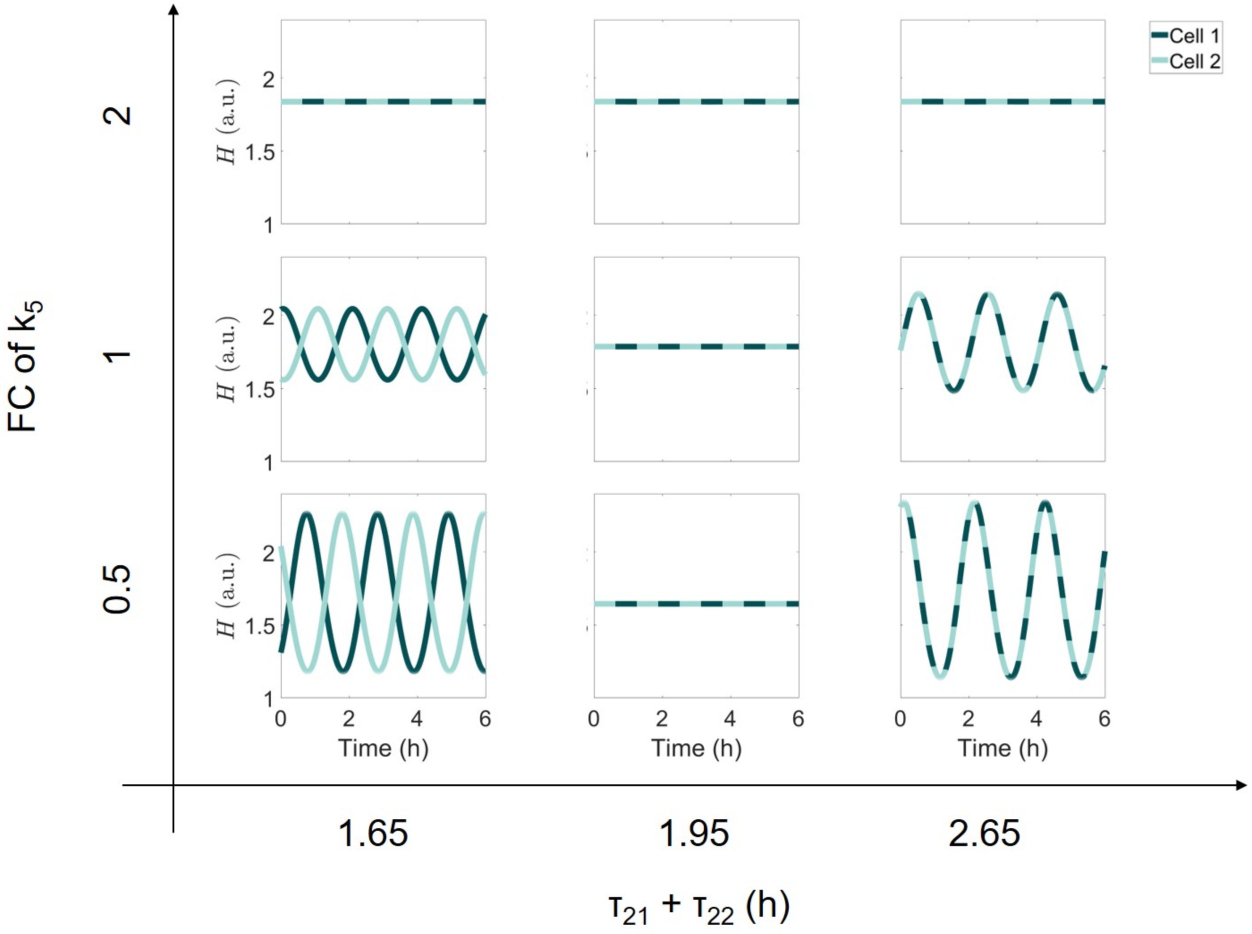


**Supplementary Figure 3 Simulations show oscillatory and sustained dynamics with respect to the total coupling delay $\tau_{21} + \tau_{22}$ and the parameter $\mathbf{k_5}$.**

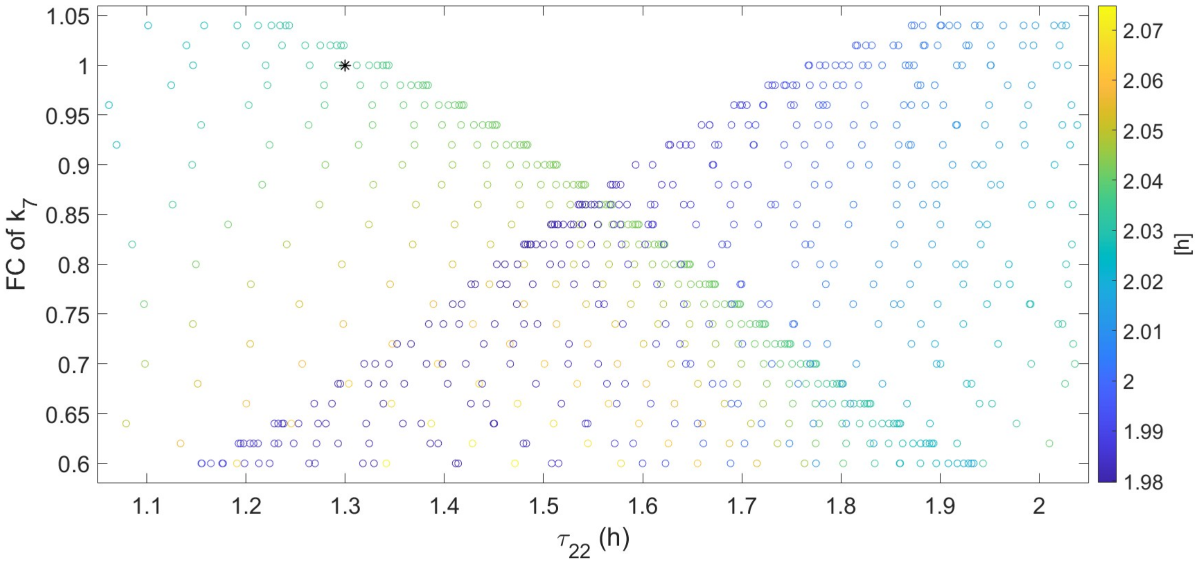


**Supplementary Figure 4 Impact of intercellular coupling delay and inverse of coupling strength on period of different periodic orbits.** Point of reference is at $\tau_{22} = 1.3$ h and $k_7 = 1$ FC marked by a star.
